# Comparative CNS Pharmacology of the Bruton’s Tyrosine Kinase (BTK) Inhibitor Tolebrutinib Versus Other BTK Inhibitor Candidates for Treating Multiple Sclerosis

**DOI:** 10.1101/2024.03.25.586667

**Authors:** Timothy J. Turner, Priscilla Brun, Ross Gruber, Dimitry Ofengeim

**Author notes:** Currently employed by Takeda (Ross Gruber).

## Abstract

Tolebrutinib is a covalent BTK inhibitor designed and selected for potency and CNS exposure to optimize impact on BTK-dependent signaling in CNS-resident cells. We applied a translational approach to evaluate three BTK inhibitors in Phase 3 clinical development in MS with respect to their relative potency to block BTK-dependent signaling and exposure in the CNS, employing *in vitro* kinase, cellular activation assays, and pharmacokinetic sampling of cerebrospinal fluid (CSF) in the non-human primate cynomolgus to estimate the ability of these candidates (evobrutinib, fenebrutinib, and tolebrutinib) to block BTK-dependent signaling inside the CNS. *In vitro* kinase assays demonstrated that tolebrutinib reacted with BTK 65-times faster than evobrutinib, while fenebrutinib, a classical reversible antagonist with a K_i_ value of 4.7 nM and slow off-rate (1.54 x 10^−5^ s^-1^), also had an association rate 1760-fold slower (3.28 x 10^3^ M^-1^ * s^-1^). Estimates of cellular potency were largely consistent with the *in vitro* kinase assays, with an estimated IC50 of 0.7 nM for tolebrutinib against 34.5 nM for evobrutinib and 2.9 nM for fenebrutinib. We then observed that evobrutinib, fenebrutinib, and tolebrutinib achieved similar levels of exposure in non-human primate CSF after oral doses of 10 mg/kg. However, tolebrutinib CSF exposure (4.8 ng/mL) (kp,uu CSF=0.40) exceeded the IC90 (the estimated concentration inhibiting 90% of kinase activity) value, while evobrutinib (3.2 ng/mL) (kp,uu CSF=0.13) and fenebrutinib (12.9 ng/mL) (kp,uu CSF=0.15) failed to reach the estimated IC90 values. We conclude that tolebrutinib is the only candidate that attained relevant CSF exposure in non-human primates.

**DISCLOSURES:** **TT, PB, DO:** Employees of Sanofi (may hold shares and/or stock options in the company).

**RG:** none.

## Introduction

There are at least 20 approved disease-modifying therapies (DMTs) to treat multiple sclerosis (MS) currently available, in large measure due to the use of the radiographic measure of contrast-enhancing T1 lesions on magnetic resonance imaging (MRI), a predictive biomarker for clinical outcomes in Phase 3 registrational trials[1, 2]. Current “high-efficacy” therapies, such as monoclonal antibodies targeting CD20, a cell surface marker expressed primarily by B lymphocytes, have largely eliminated acute focal inflammation that drives relapses and CNS lesions as observed on MRI [3, 4]. In spite of this remarkable achievement, even the most effective DMTs fall short of achieving a complete therapeutic response, as disability accumulation proceeds in most people with MS (pwMS) even in the absence of relapses and new lesion formation, a phenomenon known as Progression Independent of Relapse Activity (PIRA)[5].

The inability of conventional immunomodulatory therapies to achieve complete and comprehensive remission of disease activity reveals that adaptive immunity driven by peripheral lymphocytes has reached a ceiling in the quest for complete remission, strongly suggesting that other pathological processes resistant to conventional DMTs are driving the residual disability accumulation[6–8]. Thus, investigators have sought alternative hypotheses to account for this discrepancy in a quest to develop next-generation DMTs. Because existing DMTs act predominantly or exclusively on adaptive immunity in the periphery, attention has turned to pathological processes that are compartmentalized behind the Blood-Brain Barrier (BBB). In particular, microglia have come into focus as potential culprits in the resistant disease activity[9–12]. Microglia comprise more than 10% of the total cells in the CNS and are the chief tissue-resident cells of immunity. Embryologically, microglia represent a branch of the innate immune system derived from the yolk sac early in development distinct from neural crest derived myeloid cells that originate from the primordial liver. Microglia serve as the “first-responders” to immunological threats to the CNS, and are able to recruit lymphocytes from the periphery to mount a robust immune response to immunological insult. Because microglia reside behind the BBB, developing drugs to modulate microglial activity is challenging due to the strict regulation of entry of exogenous molecules across the BBB and into the parenchyma.

In our efforts to develop a next-generation DMT that can directly modulate neuroinflammation, we have focused on Bruton’s Tyrosine Kinase (BTK) as an attractive candidate target[13]. BTK is a non-receptor tyrosine kinase expressed in most hematopoietic cell lineages, including B lymphocytes and myeloid cell types. BTK mediates signalling from certain cell surface receptors (e.g., BCR, FcγR, TLR) to downstream elements crucial to immune cell function. Because BTK is essential for coupling extracellular stimuli to activation of an immune response in both adaptive immunity (B lymphocytes) and myeloid phagocytic cells (CNS microglia and bone marrow-derived monocytes/macrophages), inhibiting BTK presents the opportunity to blunt adaptive and innate immune responses linked to MS pathology. However, in order to effectively modulate microglial activity, candidate DMTs need to evade the multiple mechanisms that serve to exclude xenobiotics from entering the parenchymal sanctuary domain[14–16].

Modulating BTK signalling within the CNS is a central tenet of the tolebrutinib development program. Tolebrutinib is a small molecule that irreversibly inactivates BTK by a covalent reaction between its acrylamide “warhead” and the sulfhydryl sidechain of cysteine 481 in the ATP binding pocket, rendering the enzyme catalytically inert[17]. Tolebrutinib was specifically designed and selected for nanomolar potency and the ability to cross the BBB. Preclinical data, along with CSF exposure data obtained in Phase 1 trials in healthy human volunteers has demonstrated that tolebrutinib achieves sufficient exposure to engage and inactivate BTK within the CNS[18].

Within a highly competitive space for developing more effective DMTs to treat the unmet medical need in MS, numerous competitors have advanced BTK inhibitors as investigational candidates. In a Phase 2b Proof of Concept trial[19], we demonstrated that tolebrutinib was able to effectively suppress new lesion formation within 12 weeks of treatment, an effect that is a well-established predictive biomarker for clinical efficacy in Phase 3 registrational trials in relapsing forms of MS. However, our working hypothesis is that in order to diminish disability accumulation refractory to conventional DMT immunomodulators, candidate therapies need to cross the BBB to a degree sufficient to curtail pathological processes within the CNS. Unfortunately, there is little clarity as to which of the several investigational BTK inhibitors possess the appropriate attributes of potency and CNS penetrance to modulate BTK within the brain and spinal cord. Accordingly, we sought to characterize the relative ability of three candidate BTK inhibitors currently in Phase 3 development (evobrutinib, fenebrutinib, tolebrutinib) to penetrate the blood-brain barrier with sufficient exposure to inhibit the intended target.

To achieve this goal, we took a translational approach to characterize the three candidates. We compared *in silico* assessments based on Lipinski’s Rule of Five[20], *in vitro* kinase assays, cellular assays of apparent potency, and pharmacokinetic assessment of exposure of the candidates in plasma and CSF of the non-human primate cynomolgus (Macaca fascicularis) to predict the rank order of potency to inhibit BTK signalling in CNS-resident cells in human.

## Methods and Materials

### *In silico* prediction of CNS penetrance

Multiparameter optimization (MPO) scores were derived from computationally accessible physicochemical descriptors. All values for clogP, logD (pH 7.4), and polar surface area (PSA) were computed using ACDlabs (version 12). The number of H-donors was obtained from RDKit (version 2021.09.1), while the pKa value for the most basic center was computed using MoKa (version 2.6.6).

### Biochemical kinase assays

BTK activity was assessed by a third-party vendor (Nanosyn, Santa Clara, CA) using an assay mixture containing defined concentrations of inhibitor, 16 mM ATP, along with artificial, fluorescently-labeled substrate (FAMGEEPLYWSFPAKKK-NH2). The reaction was initiated by adding BTK (0.5 nM), and the reaction was followed for 3 hrs. at room temperature. A 12-point concentration response curve was constructed using the standard BTK kinase assay conditions. Phosphotransferase activity was monitored optically using a microfluidic Caliper system. Inhibitors were provided to the vendor in a blinded fashion, diluted in 100% DMSO using serial 3-fold dilution steps. Final compound concentrations in the assay ranged from 10 μM to 0.0565 nM. Compounds were tested in a single well for each dilution step, and the final concentration of DMSO was kept at 1% for all wells. A reference compound, staurosporine, was tested in an identical manner.

### Cellular activation assays

Ramos cell (an immortalized human B cell line) expression of CD69 was used to evaluate cellular activity linked to BTK signaling[21, 22]. Defined concentrations of inhibitor were obtained via twelve serial 3-fold dilution steps ranging between 6 μM and 33.8 pM. B cell receptor was stimulated by adding goat α-Human IgM F(ab’)2 at a final concentration of 20 μg/ml. The activated Ramos cells were incubated overnight (18 h) in 5% CO2 at 37°C followed by staining for flow cytometry (BD LSR2) and analysis (FlowJo and Graphpad Prism).

### Non-Human Primate (NHP) PK Studies

Three healthy male animals were used in a crossover study conducted by a third-party vendor (Biomere, Worcester, MA). On Days 1-4, animals received a single daily oral dose of the test article at 3 or 10 mg/kg. The dose volume was 5 mL/kg. Following a 7-day washout period, the same three animals received a single daily oral dose of a second test article at 3 or 10 mg/kg. This sequence was repeated a third time for the final test article. Animal health checks were performed at least twice daily, in which all animals were checked for general health. Body weights were recorded prior to dosing on Day -1 or Day 1. CSF samples were collected from an indwelling intrathecal catheter accessed via subcutaneous port using sterile technique. Approximately 180 μL of fluid (saline lock) was removed prior to CSF sample collection at 1-, 2-, 4- and 8 hours post administration. All CSF samples were stored at -80ºC prior to analysis. Blood samples (∼1 mL) were collected from an appropriate peripheral vein at 0.5-, 1-, 2-, 4-, 8-, and 24-hours post administration. Whole blood was collected into K_2_EDTA tubes and placed on wet ice until processed to plasma, then stored at -80ºC for analysis.

## RESULTS

### *In silico* models predicted relative CNS permeability

Medicinal chemists have developed empirical *in silico* tools based on first principles to predict the ability of a given molecular structure to penetrate the BBB[20]. While these tools are not perfect, they provide the opportunity to guide drug design to help target pathways inside the CNS. Lipinski’s Rule of Five uses molecular attributes of molecular weight, polarity/polar surface area, partitioning between octanol and water, ionizability, and the number of hydrogen bond partners. We applied these tools to several of the relevant structures in development to derive a composite score (known as Multiple Parametric Optimization score, or MPO) used to predict the relative ability of these compounds to distribute into the CNS (Table 1). Of the six candidates evaluated, tolebrutinib ranked highest on MPO scoring.

**Table 1.** Structure-activity relationships predict CNS penetrance. The Multiple Parametric Optimization score (MPO) is an *in silico* tool to estimate the ability of a molecule to penetrate the blood-brain barrier on the basis of its chemical properties, including molecular weight (MW), calculated partition coefficient (cLogP), ionizable distribution coefficient (LogD), the number of hydrogen bond donor (H-Donors), the polar surface area (PSA), and the acid dissociation constant (pKa).

| Test article | MW | cLogP | LogD (pH 7.4) | H-Donors | PSA | pKa | CNS MPO |
| --- | --- | --- | --- | --- | --- | --- | --- |
| Tolebrutinib | 455.5 | 2.69 | 2.53 | 1 | 92.0 | 6.55 | 4.73 |
| Ibrutinib | 440.5 | 2.92 | 2.92 | 1 | 99.2 | 3.95 | 4.41 |
| Evobrutinib | 429.5 | 3.19 | 3.19 | 2 | 93.4 | 6.42 | 4.20 |
| Fenebrutinib | 664.8 | 1.59 | 1.57 | 2 | 119.3 | 7.55 | 3.52 |
| BIIB 091 | 542.6 | 0.20 | 0.19 | 2 | 127.9 | 6.75 | 3.50 |
| Remibrutinib | 507.5 | 3.29 | 3.29 | 2 | 110.4 | 5.65 | 3.03 |

### Inhibitor potency in a cell-free kinase assay

Three candidates, evobrutinib, fenebrutinib, and tolebrutinib were selected for further pharmacological characterization based on their status as Phase 3 development candidates to treat pwMS. The candidate compounds were supplied to a vendor in a blinded fashion, and a standard two-step screening process was used. The first step was to screen a broad kinase panel of 250 protein kinases using a fixed concentration of test article at 1 μM, followed by a concentration-inhibition assay for those kinases identified in the first step as inhibition greater than 90% at 1 μM. These are pseudo-equilibrium measurements for evobrutinib and tolebrutinib as irreversible covalent inhibitors, but when conducted under identical conditions, they provide an estimate of rank-order of potency as a correlate of reaction speed at a given concentration. These measurements indicated that tolebrutinib was the most potent of the three (IC50 = 0.676 ± 0.7 nM), which was 50-fold more potent than evobrutinib (IC50 = 33.5 nM) and 9.3-fold more potent than fenebrutinib (IC50 = 6.21 nM) under these conditions.

### Kinetic approach to measuring relative potency

We compared the reaction rates (as opposed to the pseudo-equilibrium measurements provided in the screening assays) by measuring the rate of kinase inhibition at pharmacologically-relevant concentrations of the inhibitors. The two irreversible inhibitors were assumed to have a dissociation rate from the enzyme (i.e., k_off_) of zero, such that the reaction rate could be modelled by a two-step process of binding followed by irreversible reaction of the warhead with the sulfhydryl side chain of C481 of the BTK enzyme. Using a microfluidic sampling method, we determined a parameter of “Kinact/KI” which is a conventional means of estimating the relative reaction rates for these inhibitors[23]. Fenebrutinib is a classical reversible inhibitor that was designed to interact with and stabilize the inactive conformation of the kinase[24]. As such, the dissociation kinetics are relatively slow, reported as a dissociation rate of 1.52 x 10^−5^ s^-1^. Taken together, we estimated that tolebrutinib reacts with BTK at a rate 64-times faster than evobrutinib (4.37 μM^-1^ s^-1^ vs. 0.068 μM^-1^ s^-1^), and 1768-times faster than the fenebrutinib association rate constant (0.00245 μM^-1^ s^-1^) (fig. 1). This illustrated a thermodynamic liability of fenebrutinib in that slow dissociation rate constants dictate that the corresponding first-order rate constant for association is also quite slow, such that even with b.i.d. dosing, the approach to steady-state inhibition is predicted to be protracted and incomplete in the CNS.

**Fig 1.**
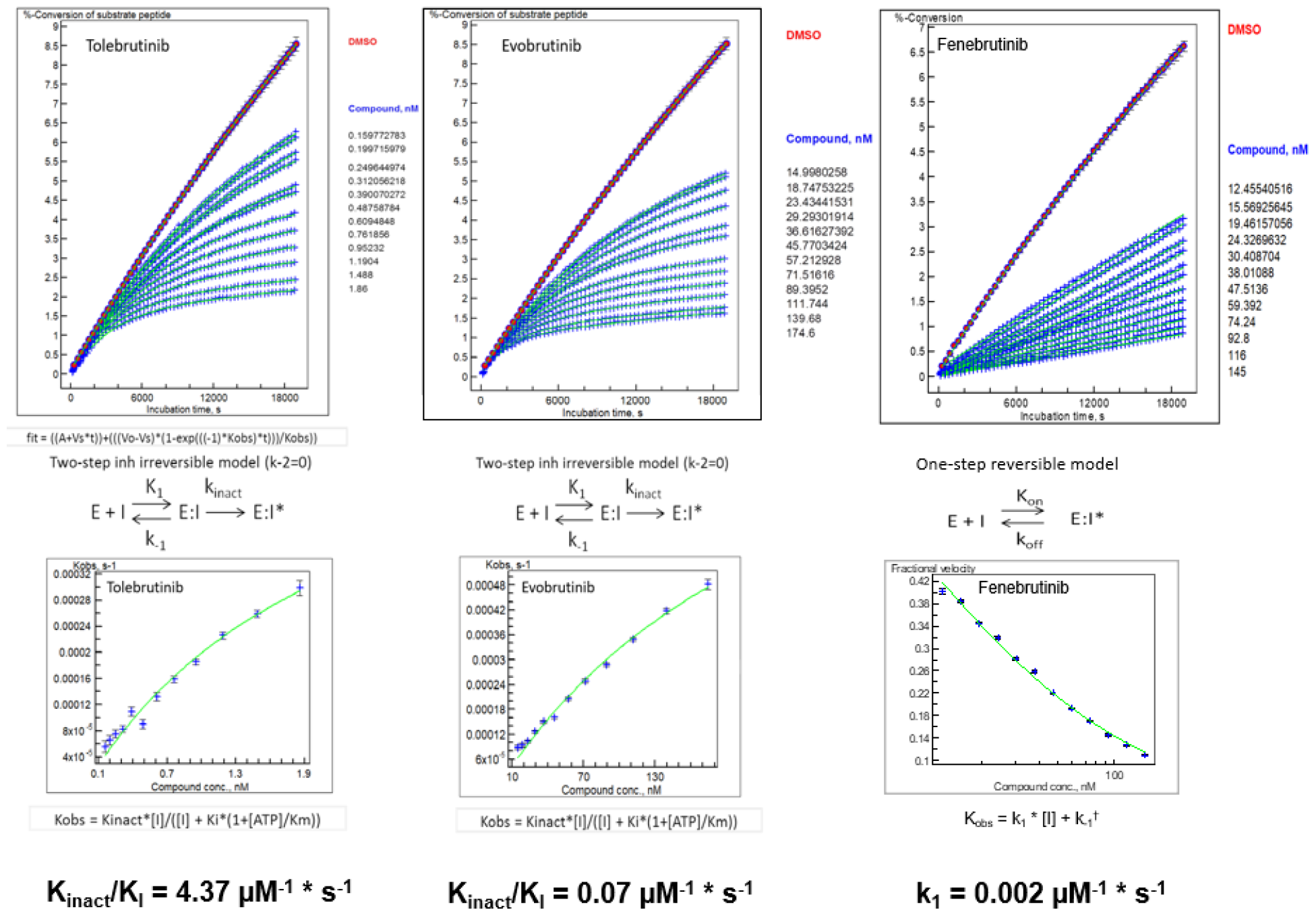
Microfluidic kinetic assay to measure BTK phosphotransferase activity as a function of inhibitor concentrations. (a-c) The observed rate of phosphokinase activity was estimated using a microfluidic assay coupled to electrophorectic separation of peptide substrate from phosphopeptide product. Sixty samples per well were collected at precisely determined times (∼ 5 minute-intervals) over the course of ∼315 minutes. Each data point represents the ratio of conversion of pseudosubstrate to product as a function of time from an individual well containing 1% DMSO (uninhibited enzyme reaction rate, n=6), an enzyme-free reaction to establish 100% inhibition (n=6), or the indicated concentration of inhibitor (n=1 for each of 12 concentrations) displayed along the right margin for each panel. (d-f) The family of curves for each analyte were fitted using three models (one-step reversible, one-step irreversible, two-step irreversible), and the best-fit model was used to determine Kobs for each article. These data were then transformed using the best-fit model to determine Kinact/Ki (tolebrutinib, evobrutinib) or Ki (fenebrutinib) to estimate the relative reaction rates for each test article.

### *In vitro* cellular potency

Biochemical potency measurements are a useful first step in understanding the pharmacology of these kinase inhibitors. However, as BTK is an intracellular non-receptor tyrosine kinase, the ability of such inhibitors to block intracellular signalling through BTK-dependent pathways is often limited by the ability of the inhibitors to gain access to the cytoplasmic compartment where the enzyme resides. Further, the same mechanisms that exclude xenobiotics from the CNS operate to protect the cytoplasm.

The relationship between extracellular concentrations of drug and the inhibition of intracellular signalling in cells of interest was investigated using the Ramos cell line. This is an immortalized cell line derived from human lymphoma that retains many functional properties of non-transformed B lymphocytes that is commonly used as a model of B cell biology. Cell surface expression of CD69, a commonly used marker of B cell activation, was used to evaluate cellular activity linked to BTK signaling[25]. Defined concentrations of the three BTK inhibitors were obtained via twelve serial 3-fold dilution steps ranging between 6 μM and 33.8 pM. B cell receptor was stimulated by adding goat α-Human IgM F(ab’)2 at a final concentration of 20 μg/ml. The stimulated Ramos cells were incubated overnight (18 h) in 5% CO2 at 37°C followed by staining for flow cytometry. A normal concentration-response curve was obtained for each of the three inhibitors, with an apparent IC50 value of 3.62, 19.8, and 89 nM for tolebrutinib, fenebrutinib, and evobrutinib, respectively (fig. 2). This represented a relative potency for tolebrutinib of 5.4-fold over fenebrutinib and 22-fold over evobrutinib. The apparent decreased potency of all three inhibitors relative to the biochemical assay likely reflects the concentration gradient between the extracellular and intracellular compartments created by various transporters (e.g., ABC and SLC transporters) as predicted.

**Fig 2.**
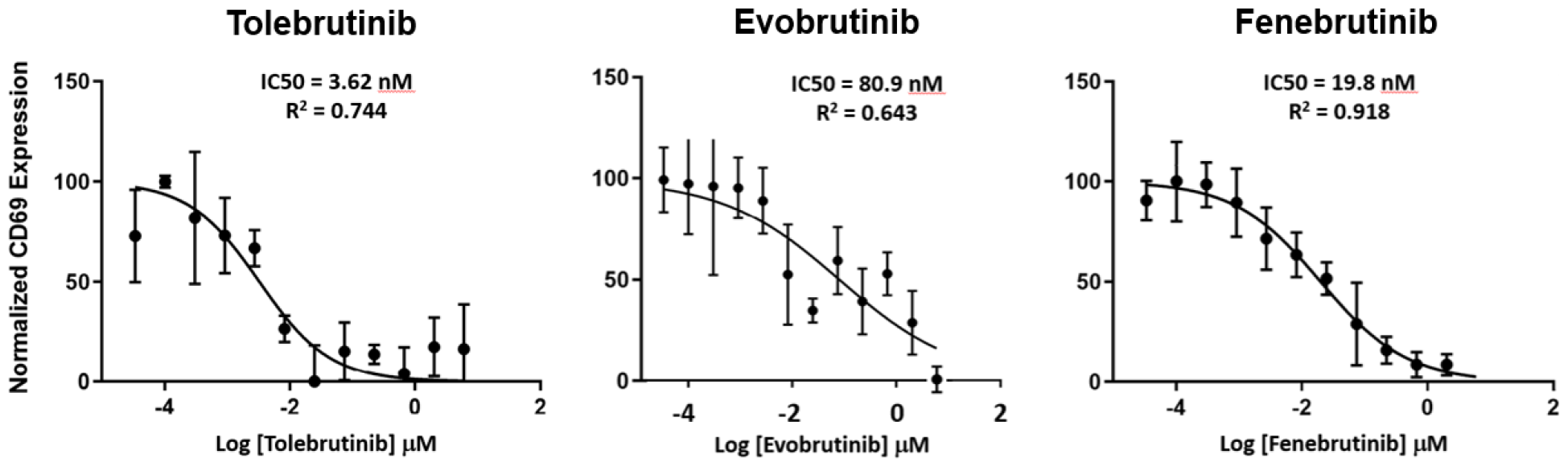
In vitro cellular potency in Ramos B-cell line. CD69 expression was used to evaluate cellular activation mediated by FcγR signalling through the BTK pathway. Defined concentrations of inhibitor were obtained via twelve serial 3-fold dilution steps ranging between 6 μM and 33.8 pM. B cell receptor was stimulated by adding goat α-Human IgM F(ab’)2 at a final concentration of 20 μg/ml. The activated Ramos cells were incubated overnight (18 h) in 5% CO2 at 37°C followed by staining for flow cytometry and analysis. Data represents the mean relative CD69 expression from triplicate measurements. Smooth curves describing the best fit to a logit model, and estimated IC50 values were generated using GraphPad.

### Tolebrutinib exposure exceeded the IC90 value in the CNS of NHPs

The foundational principle of tolebrutinib development as a next-generation therapy to treat MS is that it must achieve exposures in the CNS that will engage target and block BTK-dependent signalling within the brain and spinal cord. We conducted a pharmacokinetic study in the non-human primate cynomolgus (Macaca fascicularis) to determine the relationship between plasma and CSF exposure for the three candidates. These measurements were conducted in a blinded fashion (i.e., the test articles were provided to the vendor identified by a code known only to the sponsor) under identical conditions, using a crossover design with the same three animals and the same oral dose (10 mg/kg/day) over three consecutive days. The goal was to determine the extent to which each inhibitor achieved CSF exposure relative to the cellular potency established in the experiments reported here.

Three healthy male animals were used in a crossover study conducted by a third-party vendor (Biomere, Worcester, MA). On Days 1-4, animals received a single daily oral dose of the test article at 10 mg/kg, administered by gavage at a volume of 5 mL/kg. Following a 7-day washout period, the same three animals received a single daily oral dose of a second test article at 10 mg/kg. This sequence was repeated a third time for the final test article.

The pharmacokinetic plasma profiles of the three candidates were comparable on the basis of clearance/elimination, with a terminal t_1/2_ for elimination of ∼4 hrs for tolebrutinib and evobrutinib, with a slightly prolonged t_1/2_ for fenebrutinib as reported previously. C_max_ was greatest for fenebrutinib (553 ng/mL or 853 nM at t = 0.5 hr), followed by evobrutinib (287 ng/mL or 669 nM at t = 0.5 hr) and tolebrutinib (103 ng/mL or 226 nM at t = 1.0 hr) (fig. 3A). Given that the same dose was used for all three inhibitors, the differences in C_max_ likely reflect the extent of metabolism of each molecule, as these molecules are known to be extensively metabolized.

**Figure 3.**
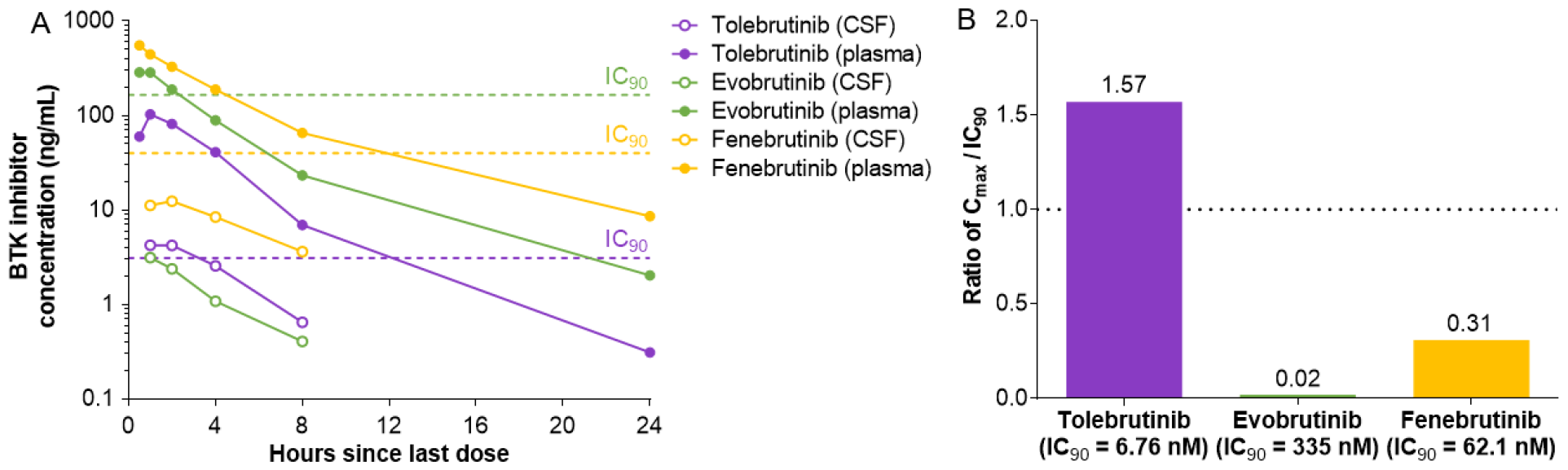
CNS Exposure of three BTK inhibitors in Non-Human Primates. Three healthy male animals were used in a crossover study conducted by a third-party vendor (Biomere, Worcester, MA). On Days 1-4, animals received a single daily oral dose of one of three test articles at 3 or 10 mg/kg that were supplied using a blinding code such that the identity of the substances was unknown to the vendor. The dose was delivered by oral gavage in a volume of 5 mL/kg. Following a 7-day washout period, the same three animals received a single daily oral dose of a second test article at 3 or 10 mg/kg. This sequence was repeated a third time for the final test article. Data presented for the 10 mg/kg dose. A) The plasma (filled symbols) and CSF (open symbols) concentrations of test article (indicated in the legend) are plotted as a function of time post-administration. The dashed horizontal lines represent the IC90 value for each test article as determined above (see Table 2). Data represents the mean values from the three animals. B) The maximal CSF exposure observed is plotted relative to the in vitro IC90 for each test article to estimate the extent to which the test article achieved bioactive exposures. At the dose of 10 mg/kg, only tolebrutinib exceeded the estimated IC90 value (1.57-fold) while evobrutinib (0.02-fold) and fenebrutinib (0.31-fold) failed to reach their respective IC90 values.

CSF exposure for each of the inhibitors was detected at the earliest time point, with a slightly slower approach to C_max_ which was observed at 1 to 2 hours post-administration (fig. 3A). The ratio of inhibitor in the CSF to plasma was 0.041 for tolebrutinib, 0.020 for fenebrutinib, and 0.0109 for evobrutinib. However, because of the differences in potency for the three inhibitors, only tolebrutinib exceeded the IC90 value for inhibiting BTK kinase activity, and in fact sustained concentrations exceeding the IC90 were maintained for at least 2 hours. Fenebrutinib was able to achieve CSF exposures roughly equivalent to the IC50 (i.e., half maximal inhibition), while the combination of poor CSF penetrance and low relative potency limited evobrutinib to C_max_ exposures well below the IC50 (7.3 nM vs the IC50 value of 33.5 nM) (fig. 3B).

**Table 2.**
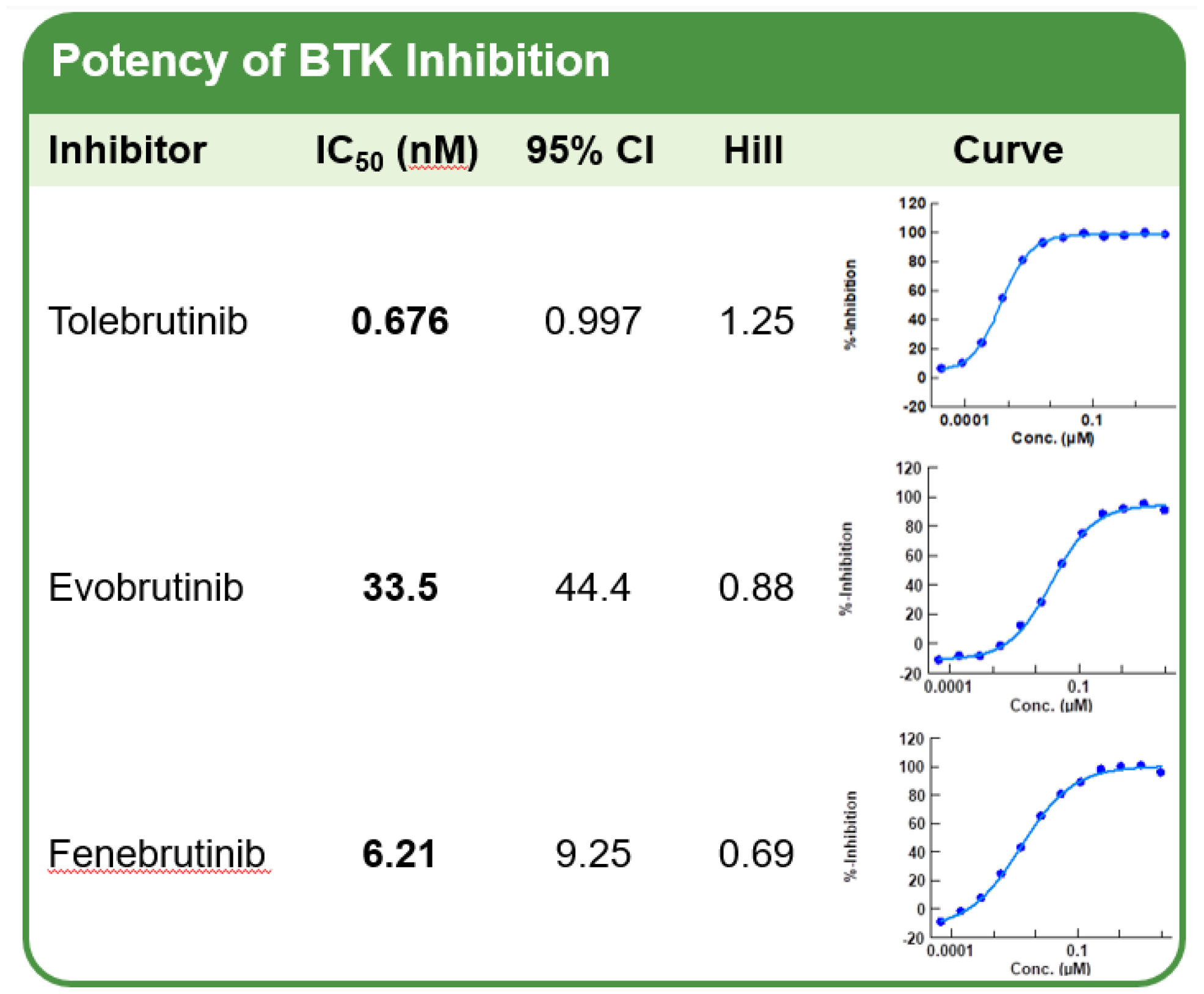
Rank-Order of Potency for Three BTK Inhibitors. Classical kinase screening assays measuring phosphokinase activity were conducted using recombinant BTK protein and a fluorescent peptide pseudosubstrate in the presence of 16 μM ATP and defined concentrations of inhibitor. The concentration response curve (right panel) was fit to a normal hyperbola adjusted using a logit calculation to estimate the IC50 and Hill slope for each test article. Results represent the average of two independent determinations.

Differences in extent of plasma protein binding determine the relative free fraction for each inhibitor. On the basis of the plasma free fractions, tolebrutinib demonstrated intrinsic CNS penetrance in cynomolgus, based on the unbound partition coefficient K_p,uu_ = 0.397, roughly 3 times higher than evobrutinib (0.131), fenebrutinib (0.147). Unbound fraction in plasma largely represents the portion of total drug not bound to plasma proteins and is related to the overall hydrophobicity.

## DISSCUSSION

The remaining unmet need for pwMS is the insidious accumulation of disability that persists even with adequate control of the focal inflammatory activity that manifests as radiographic lesions and acute relapses. Recent clinical trials of immunomodulators such as ofatumumab, ublituximab (both anti-CD20 mAbs), and the S1P modulator ponesimod have established these DMTs as being highly effective against such focal inflammation. At the same time, the impact of these agents on disability remains inadequate, as shown by Kappos et al., as roughly one in five RMS trial participants experienced one or more PIRA disability events after 96 weeks of treatment with ocrelizumab. The key question under investigation is whether compartmentalized inflammation behind the blood-brain barrier represents the limiting obstacle in preventing PIRA.

To directly modulate immune surveillance within the CNS, four conditions must be met. First, the target of interest must be expressed in cells that drive neuroinflammation that leads to pathology. Second, the drug intended to modulate the target must penetrate the blood-brain barrier. Third, the drug must attain sufficient exposure within the CNS to engage the target in a pharmacologically-meaningful manner, in principle achieving >90% trough occupancy. Fourth, engagement of the target must translate into a clinical benefit by modifying pathological processes to curtail tissue destruction and preserve function.

In the case of tolebrutinib, it was specifically designed and selected to be CNS penetrant, using the medicinal chemistry principles developed for this purpose. To translate principle to practice, we conducted the series of experiments reported here to assess the pharmacological properties of tolebrutinib to justify development of this agent in neuroinflammatory disorders like MS. Further, we sought to compare tolebrutinib with two other BTK inhibitors to benchmark tolebrutinib against these competitors. On the basis of the high potency and superior CNS penetrance, we conclude that tolebrutinib, but not evobrutinib or fenebrutinib, satisfies the first three of the four conditions described above. Validation of the fourth condition awaits completion of the pivotal trials of tolebrutinib to evaluate safety and efficacy to determine if CNS exposure translates to a clinical benefit.

Tolebrutinib was observed to be more potent but less selective than the comparators based on biochemical kinase assays. Notably, the estimated forward rates obtained in the kinetics assay (Fig. 1) revealed that the sub-nanomolar potency of tolebrutinib was also reflected in the pseudo-first order rate constant for inhibition. In particular, fenebrutinib, which binds to the inactive conformation of the enzyme, demonstrated a forward rate of binding that was ∼1780-fold slower than tolebrutinib. This observation is consistent with the different mechanistic interactions between drug and kinase, where fenebrutinib favors high selectivity over speed, while tolebrutinib favors high reaction rates while interacting with approximately seven kinases in the Tec family with a conserved binding pocket.

The consequences of these pharmacological differences are significant *in vivo*. Unlike *in vitro* experiments where the concentrations of drug are well-controlled and constant as a function of time, oral administration of a drug is a dynamic process described by the in vivo pharmacokinetics. For irreversible covalent inhibitors like tolebrutinib and evobrutinib, the reaction with the enzyme is determined by the product of the apparent forward rate constant and the duration of exposure, such that inhibition is largely proportional to the AUC. This is a significant advantage conferred upon irreversible inhibitors, as occupancy can be driven by relatively brief exposures achieved by the relatively short half-life (∼90-120 minutes). In the case of the reversible inhibitor fenebrutinib, the slow off-rate is an advantage to maintain occupancy during the inter-dose interval. However, thermodynamics requires that the slow off-rate and low nanomolar potency is accompanied by a slow first-order rate of association of the drug with kinase. This feature is a liability in a therapeutic setting, as twice daily dosing with a moderate half-life (4 to 6 hours) means that the reaction is driven primarily by the C_max_. This accounts for the relatively high plasma exposures (compared with tolebrutinib) reported for fenebrutinib, as target occupancy requires relatively high concentrations of ligand to compensate for the slow rate constant.

With regard to target engagement in the CNS, inhibiting BTK depends on achieving pharmacologically-relevant concentrations of drug for sufficient durations. Unsurprisingly, all three candidates achieved some degree of exposure in the CSF of NHPs. However, exposure *per se* is necessary, but to be relevant exposure, the ratio of exposure to potency must be sufficient to drive the binding reaction. For instance, evobrutinib achieved a C_max_ of 3.2 ng/mL or 7.5 nM, slightly less than what was observed for tolebrutinib (4.4 ng/mL, 9.6 nM). However, because the IC90 for tolebrutinib was 6.7 nM compared to 335 nM for evobrutinib, the ratio of C_max_/IC90 was 1.57 for tolebrutinib compared to 0.02 for evobrutinib. Fenebrutinib was somewhat intermediate on this measure (C_max_/IC90 = 0.31), although the potency represents an *in vitro* equilibrium measurement that is not achieved in vivo where drug concentrations are constantly changing due to absorption, distribution, metabolism and elimination.

Finally, do these observations predicting CNS exposure for the BTK inhibitors in development translate to a pharmacokinetic and pharmacodynamic effect in humans? Published data on CNS exposure of BTK inhibitors in the CNS is limited to ibrutinib[26], evobrutinib[27], and tolebrutinib[18]. Focusing on tolebrutinib, CSF exposure was determined in a Phase 1 pharmacokinetic study in healthy volunteers, using two doses (60 mg or 120 mg) administered orally in the fed state. CSF exposure estimated over a four-hour period was obtained by measuring CSF drug content at 1- and 3-hours, or 2- and 4-hours post-administration in two separate cohorts that each received two lumbar puncture procedures. CSF exposure associated with the 60 mg dose reached C_max_ of 0.87 ng/mL (1.9 nM) within 1 to 2 hours, remaining relatively stable over the four-hour observation period. This CSF exposure exceeded the IC50 value (0.70 nM) reported here for a minimum of four hours. The apparent elimination kinetics of tolebrutinib in the CSF closely resembled those reported for ibrutinib in a study with primary CNS lymphoma patients [26]. In a Phase 2 trial of evobrutinib in pwMS, a lumbar puncture obtained at a single time point two hours post-administration revealed a CSF concentration of 3.2 ng/mL (7.45 nM), an exposure approximately four-fold lower than the IC50 value (33.5 nM) reported here[27].

Methodologically, the experiments were performed by third-party vendors (the Ramos cell assays were conducted by Sanofi scientists) conducted under conditions to minimize bias favoring a given test article. Experiments were conducted so that the identity of the test articles was blinded, and the concentrations and doses used were identical with the exception of the kinetic assays where the appropriate dose ranges for each agent were determined in blinded pilot experiments. The NHP pharmacokinetic experiments were conducted in the same three ported animals by a third-party vendor. The goal was to determine, as best as possible, the relative merits and liabilities of the BTK inhibitors currently in late-stage clinical development to support rational, data-driven decisions on the development path for tolebrutinib in a highly competitive environment.

An important methodological consideration relevant to this work is the concept of thermodynamic equilibrium as it applies to covalent inhibitors. Because these chemical reactions are irreversible, true thermodynamic equilibrium is not applicable. Thus, the parameter of Kinact/Ki reported here for tolebrutinib and evobrutinib is an approximation for an equilibrium rate constant. Further, the potency values reported are also an approximation for the same reason. Because the experiments were conducted under identical conditions, the absolute value of potency is an approximation, but the rank order of potency remains relevant to interpreting the rate and extent of the inhibition expected for each agent.

In conclusion, we set out to determine whether tolebrutinib has the necessary attributes to achieve pharmacologically-relevant exposure in the CNS. Our first-principles approach started with the design and selection of tolebrutinib based on best-practice medicinal chemistry strategies, followed by pharmacological comparisons of tolebrutinib to two other BTK inhibitors in Phase 3 development to determine whether tolebrutinib is differentiated on the basis of CNS exposure. The totality of the data indicate that tolebrutinib has a combination of adequate intrinsic CNS penetrance accompanied by sub-nanomolar potency that allow CSF concentration to exceed its IC90, while fenebrutinib and evobrutinib fail to reach their respective IC90 levels in non-human primates. Thus, we have established that 1) BTK is present in cells implicated in the pathobiology of MS, 2) that tolebrutinib has excellent CNS penetrance, and 3) that the CSF exposure achieved by tolebrutinib exceeds its IC90 in non-human primates. What remains to be determined, 4) that tolebrutinib exposure translates a clinical benefit with respect to modulating neuroinflammation, awaits confirmation in the Phase 3 clinical trials.

